# Identification of ribosomal protein eL21 as a novel externalized protein and a potential target in triple negative breast cancer

**DOI:** 10.1101/2025.03.25.645355

**Authors:** Lucie Arnould, Nina Radosevic-Robin, Jérémy Néri, Frédérique Penault-Llorca, Jean-Jacques Diaz, Marie Alexandra Albaret, Frédéric Catez

## Abstract

Proteins normally localized in the intracellular compartments of healthy cells, have been observed at the surface of cancer cells, despite not having transmembrane domain or secretion signals. The unexpected localization of these externalized proteins is likely to reflect currently unknown functions. Their presence on the cell surface presents a unique opportunity for cancer therapy, enabling the development of antibody- or peptide-based strategies. In this study, we demonstrate the presence of an extracellular form of the ribosomal protein L21 (eL21) in triple negative breast cancer cells, using multiple complementary approaches and a broad set of antibodies. Importantly, we show that anti-eL21 antibodies induce a potent and immediate anti-proliferative effect characterized by cell cycle arrest and apoptosis. Our findings, uncover eL21 as novel ribosomal protein exhibiting extra-ribosomal functions. Finally, our study identifies eL21 as a promising therapeutic target in triple negative breast cancer.

## Introduction

Triple-negative breast cancers (TNBC) account for 15 to 20 % of breast cancers and 83 % of deaths related to breast cancer [1], [2]. TNBC cells lack expression of estrogen and progesterone receptors and do not display amplification of the human epidermal growth factor receptor 2 (HER2) gene [1]. TNBC is more proliferative, aggressive and metastatic, than other breast cancer subtypes and present a higher risk of recurrence [3], [4]. The predominant systemic chemotherapy [5], [6], including cisplatin and anthracyclines/taxane-based regimens, shows limited efficacy due to the heterogeneity of oncogenic drivers [7], [8], [9] and the development of resistance [7], [8], [10], [11]. As a result, the median overall survival for patients is only of 12 to 18 months [4]. Given these clinical challenges, it is crucial to explore new therapeutic strategies.

Recent studies have revealed the unexpected presence of proteins at the surface of the tumor cells that are normally localized in the intracellular compartment of healthy cells. These proteins, also referred to as externalized proteins, lack transmembrane domain or signal peptide, and the anchoring and export mechanisms of the proteins to the cell surface remain unknown [12], [13]. Notably, externalized proteins may exhibit novel functions distinct from their intracellular counterparts, modulating processes such as proliferation, invasion, and apoptosis, thereby contributing to cancer progression [14], [15]. For instance, the interaction between the externalized form of glucose-regulated protein 94 (GRP94), estrogen receptor α36 (ER-α36), and HER2, on the surface, appears to active a feedback loop via cell membrane gp96 in TNBC [16], [17]. Similarly, nucleolin and ER-α36 are overexpressed in lymphoma and breast cancer cells and interact with the epidermal growth factor receptor (EGFR), which helps to enhance tumor progression [18], [19], [20], and externalized Histone 2B and cytokeratin 8 (CK8) are thought to play a role in invasion by acting as plasminogen receptors [21], [22], [23]. We previously reported the presence of an externalized form of the intermediate filament protein CK8 at the surface of colorectal and head & neck cancer cells and made the proof of concept that externalized CK8 represent an actionable target using a monoclonal anti-CK8 antibody [13], [22]. Given their accessibility and cancer-specific expression, externalized proteins are emerging as promising candidates for targeted cancer therapies.

Ribosomal protein (RP) gene expression is altered in numerous cancer such as breast, colon and prostate cancer, and these changes are thought to influence tumor cell behavior [22], [24], [25]. While altered RP expression may impact cellular translation, it can also impact other cellular processes through the moonlighting functions of RPs. Indeed, several RPs exhibit extra-ribosomal functions due to their ability to interact with non-ribosomal RNAs and proteins [26]. These extra-ribosomal activities include regulating transcription, splicing, contributing to viral infection, and interacting with key regulatory factors such as c-jun or NFkB. Additionally, RPs can modulate p53 level by binding of MDM2 [27]. Among candidate RPs with extra-ribosomal function, the ribosomal protein P0 (uL10) was recently found at the plasma membrane of breast cancer and head & neck cancer cells [28], [29]. Notably, autoantibodies directed against uL10 was found in sera of breast cancer patients, and immunization with uL10 protein delayed the onset of mammary tumors in a mouse model of age-related breast cancer [28]. Therefore, exploring the extra-ribosomal functions of RPs could reveal novel mechanisms underlying tumorigenesis and present new opportunism for developing cancer-specific therapies.

In this study, we demonstrate that ribosomal protein L21 (eL21) is unexpectedly present on the surface of TNBC cells. Notably, we show that an antibody targeting this externalized form of eL21 inhibits tumor growth, induces cell cycle arrest and triggers the intrinsic apoptotic pathway. Overall, our findings reveal that externalized eL21 is present in tumor cells and represents a promising therapeutic target for the treatment of triple-negative breast cancer.

## Materials and Methods

### Cell lines

TNBC cell lines CAL51 and Hs 578T were generously provided by Dr Thierry Dubois (Curie Institute, Paris, France). Cells were maintained in Dulbecco Minimum Essential Medium - GlutaMax^TM^ (Invitrogen, supplemented with 10 % fetal bovine serum (FBS) and 1 % of Penicillin/Streptomycin (Invitrogen) at 37 °C with 5 % CO_2_.

### Antibodies

For immunofluorescence, the primary antibodies used were: rabbit polyclonal anti-eL21 (Sigma-Aldrich, HPA047252), mouse polyclonal anti-eL21 (Abcam, ab194664), rabbit polyclonal anti-eL21 (Bethyl, A305-032A), mouse monoclonal anti-eL21 clone 2D8 (Abnova, H00006144-M03), mouse (Abcam, ab7671) and rabbit (Cell Signaling Technology, 3010S) anti-alpha 1 sodium potassium ATPase, at 1:100 dilution.

For western blot analysis, the primary antibodies used were: rabbit polyclonal anti-eL21 (Bethyl, A305-032A), rabbit polyclonal anti-ribosomal protein L29 (eL29) (Bethyl, A305-055A), rabbit polyclonal anti-ribosomal protein L5 (uL18) (Abcam, ab86863), rabbit monoclonal anti-uL10 (Abcam, ab52947), rabbit polyclonal anti-ribosomal protein S2 (uS5) (Bethyl, A303-793A) and mouse (Abcam, ab7671) anti-alpha 1 sodium potassium ATPase, at 1:1000 dilution.

For electron microscopy analysis, the primary antibodies used were: rabbit polyclonal anti-eL21 (Bethyl, A305-032A) and rabbit polyclonal anti-EGFR (Cell Signaling Technology, 2232S) at 1:100 dilution.

For microbeads binding assays, the primary antibody used was a rabbit polyclonal anti-eL21 (Bethyl, A305-032A) at 1:100 dilution.

For functional analysis, the primary antibodies used were: rabbit polyclonal anti-eL21 (Sigma-Aldrich, HPA047252) and a rabbit polyclonal IgG isotype control (R&D system, AB-105-C). The immunogen used to raise eL21 antibodies are depicted in the Supplementary Figure 1.

### Immunofluorescence

Cells were plated on glass coverslips (Knittel) in 24-well plates. Primary antibodies were revealed by goat anti-rabbit secondary antibody coupled to Alexa Fluor-488 (Life Technologies, A11034) at 1:500 dilution, goat anti-mouse secondary antibody coupled to Alexa Fluor-488 (Life Technologies, A11029) at 1: 500, goat anti-rabbit secondary antibody coupled to Alexa Fluor-555 (Life Technologies, A21428) at 1:1000 dilution and goat anti-mouse secondary antibody coupled to Alexa Fluor-555 (Life Technologies, A21424) at 1:1000 dilution. Nuclei were labeled with Hoechst (Sigma, B2261) at 1:10,000 dilution in 1X PBS. Fluoromount G (Electron Microscopy Science, 17984-25) was used for mounting coverslips on glass slides. Negative control conditions were cells incubated with secondary antibodies only. Fluorescence images were acquired using excitation wavelengths of 405 nm, 488 nm and 555 nm in confocal microscopy, on a Zeiss LSM 880 confocal microscope using a Plan Apochromat 63X immersion objective. Images were then processed using Zen® software.

#### Intracellular staining

Cells were fixed with 4 % paraformaldehyde (PFA) (Electron Microscopy Science, 15710) in 1X PBS for 30 minutes at room temperature. The cells were then permeabilized with 0.5 % Triton X-100 (Sigma-Aldrich, X100) in 1X PBS for 5 minutes and blocked in antibody buffer containing 1 % bovine serum albumin (BSA) (Sigma-Aldrich, A7030), 2.2 % sodium chloride (NaCl) (Sigma-Aldrich, S3014), and 0.5 % Tween 20 (Sigma-Aldrich, P1379) for 2 hours. Next, the cells were incubated with the primary antibody diluted in the antibody buffer solution for 1 hour at room temperature, followed by incubation with secondary antibodies conjugated to Alexa Fluor 488 and Alexa Fluor 555 for 45 minutes at room temperature in the dark. Finally, nuclear staining was performed for 5 minutes at room temperature in the dark. All washing steps were done with 1X PBS containing 10 mM glycine (Sigma-Aldrich, G8898).

#### Cell surface staining

Cells were fixed with 4 % PFA solution for 2 minutes and 30 seconds at room temperature. The cells were then incubated with the primary antibody diluted in 1X PBS containing 3 % BSA for 2 hours at room temperature, followed by incubation with secondary antibodies conjugated to Alexa Fluor 488 and Alexa Fluor 555 for 45 minutes at room temperature in the dark. Finally, nuclear staining was performed for 5 minutes at room temperature in the dark. All washing steps were carried out with 1X PBS.

### Subcellular fractionation

Subcellular fractionation and plasma membrane preparation were performed using a Plasma Membrane Protein Extraction kit (Abcam, ab65400) according to manufacturer guidelines.

### Western Blot

The cells were then lysed by manual scraping to extract proteins using 2X Laemmli lysis buffer, containing 0.125 M Tris-HCl at pH 6.8, 2 % sodium dodecyl sulfate (SDS) (Fisher Scientific, BP1311-1), 200 mM dithiothreitol (DTT) (Sigma-Aldrich, D8255), 20 % glycerol (Euromedex, 50405), and distilled water. Protein lysates were then heated at 95 °C for 10 minutes and proteins were quantified using the trichloroacetic acid (TCA) assay (Sigma-Aldrich, T0699), reading absorbance at 570 nm (Tecan, Life Sciences). Proteins were separated by SDS-PAGE with migration buffer containing 25 mM Tris, 192 mM glycine, and 0.1 % SDS at pH 8.3 (Bio-Rad, 161-0772) for 1 hour at 140 V at room temperature. Transfer was then carried out for 1 hour at 100 V at room temperature with transfer buffer containing 25 mM Tris and 192 mM glycine at pH 8.3 (Bio-Rad, 161-0771) and nitrocellulose membranes (Amersham Protan Premium 0.45 NC, 10600003). The membranes were then blocked in a saline-Tris buffer (TBS) solution (Fisher Scientific, 3769684) containing 1 % Tween-20 and 5 % milk for 1 hour at room temperature with agitation. The membranes were then incubated with primary antibodies diluted in TBS with 1 % Tween-20 and 5 % milk, with agitation, for 1 hour at room temperature or overnight at 4 °C. The membranes were washed with TBS containing 1 % Tween-20 three times for 10 minutes at room temperature with agitation. Secondary antibodies were incubated for 1 hour at room temperature with agitation. The membranes were washed with TBS containing 1 % Tween-20 three times for 10 minutes at room temperature with agitation. Finally, detection was performed by chemiluminescence, incubating the membranes with Clarity™ Western ECL Substrate (Bio-Rad, 170-5061) for 5 minutes at room temperature, followed by imaging with a cold-sensitive camera (ChemiDoc, Western Blot Imaging, Bio-Rad). Images were processed using ImageLab® software.

### Electron microscopy

Cells were fixed at 37 °C in 4 % PFA complemented with 0.2 % glutaraldehyde. Cells were washed three times in cacodylate 0.2 M saccharose 0.4 M for 1 hour at 4 °C, dehydrated though a series washes with 30, 50, 70 % ethanol maintained prealably at 4 °C for 5 minutes, infiltrated with London Resin White (LRWhite, EMS, France) using 1:1 LRWhite and 4 °C absolute ethanol for 60 minutes followed by pure LRWhite at 4 °C for three periods of 60 minutes each, then embedded in pure LRWhite in gelatine capsules for polymerization at 50 °C for 48 hours. Ultrathin sections (approximately 70 nm thick) were cut on a UC7 (Leica) ultra-microtome, mounted on 200 mesh nickel grids coated with 1:1,000 polylysine, and stabilized for 1 day at room temperature. Immunogold labelling was performed by flotation the grids on drops of reactive media. Nonspecific sites were coated with 1 % BSA and 1 % normal goat serum in 50 mM Tris-HCl, pH 7.4 for 20 minutes at room temperature. Thereafter, incubation was carried out overnight at 4 °C in wet chamber with primary antibody (Bethyl, A305-032A). Sections were successively washed three times in 50 mM Tris-HCl, pH 7.4 and pH 8.2 at room temperature. They were incubated in a wet chamber for 45 minutes at room temperature in 1 % BSA, 50 mM Tris-HCl, pH 8.2 for 20 minutes at room temperature, labeled with gold conjugated secondary antibody (Aurion). Sections were successively washed three times in 50 Mm Tris-HCl pH 8.2 and pH 7.4 and three times infiltrated distilled water. The immunocomplex was fixed by a wash in glutaraldehyde 4 % for 3 minutes. Sections were stained with 0.5 % uranyl acetate in ethanol 50 % for 5 minutes in darkness and observed with a transmission electron microscope JEOL 1400JEM (Tokyo, Japan) operating at 80 kV equipped with a camera Orius 600 Gatan and Digital Micrograph.

### Microbeads binding assay

Cells were plated on glass coverslips in 24-well plates. Cells were fixed with 4 % PFA in 1X PBS for 2 minutes and 30 seconds at room temperature. Saturation was carried out in 1X PBS containing 3 % BSA for 30 minutes at room temperature. Cells were incubated with the primary antibody diluted in 1X PBS containing 3 % BSA for 2 hours at room temperature, followed by incubation with protein G microbeads for 45 minutes at room temperature. Finally, nuclear staining was performed for 5 minutes at room temperature in the dark. All washing steps were performed with 1X PBS. Primary antibodies were detected using protein G microbeads (Thermo Fisher Scientific, 1004D) at a 1:1,000 dilution. Nuclei were stained with Hoechst (Sigma-Aldrich, B2261) at a 1:10,000 dilution in 1X PBS. Negative control conditions consisted of cells incubated with protein G microbeads only. Images were acquired by phase contrast imaging on a Zeiss AxioVert microscope using a 40X objective. The images were then processed using Zen® software. Images contrast was adjusted to enhance beads visualization. Bead counting via High-Content Screening (HCS) was performed on a Revvity Operetta-CLS system with a 20X NA 0.4 objective. Image acquisition was done using Harmony software (Revvity). Image analysis and nucleus counting were performed with Columbus software (Revvity).

### EdU incorporation assay

Cells were placed in 96-well plates and incubated for 96 hours with or without Staurosporine (Sigma-Aldrich, S6942), with a rabbit anti-eL21 polyclonal antibody, or with a rabbit polyclonal IgG isotype control. Each condition was performed in triplicate. EdU at 1 mM (Jena Bioscience, CLK-N001) was added to the appropriate cell culture medium for 60 minutes at 37 °C and 5 % CO_2_. The cells were fixed with 4 % PFA in 1X PBS for 5 minutes at room temperature. Cells were permeabilized with 0.5 % Triton X-100 in 1X PBS for 20 minutes. The cells were then incubated in 1X PBS containing 2 mM CuSO_4_·H_2_O (Sigma-Aldrich, 12849), 4 *µ*M Sulfo-Cy^3^-Azide (Lumiprobe, B1330), and 20 mg/mL ascorbic acid (Sigma-Aldrich, A4544) for 30 minutes at room temperature, in the dark and under paraffin to protect from oxidation. Finally, nuclear staining was performed with Hoechst for 5 minutes at room temperature in the dark. All washing steps were performed with 1X PBS containing 20 mM glycine.

### Real-time analysis of cell confluence and apoptosis

For each condition, cells were seeded in flat-bottom 96-well plates. The cells were untreated, treated with Staurosporine, treated with the rabbit anti-eL21 polyclonal antibody, and treated with a rabbit polyclonal IgG isotype control. Each condition was performed in triplicate. Images were taken every two hours for 120 hours to analyze cell confluence and for 96 hours for caspase-dependent apoptosis, at 4X magnification, using the IncuCyte ZOOM® Live Content Imaging System® (Sartorius). Cell confluency was determined in each image by measuring the area occupied by the cell, relative to the total surface of the field of view. For Caspase 3/7 activation analysis, cells were incubated with the caspase-3/7 apoptosis reagent (Sartorius) at a 1:1,000 dilution per well. The total fluorescence signal was integrated and normalized to cell confluency.

### Spheroids formation

For each condition, cells were seeded in 96-well ULA round-bottom plates for spheroids (Perkin Elmer, 6055330). The cells were untreated, treated with Doxorubicin (Accord Pharma), or treated with the rabbit anti-eL21 polyclonal antibody. Each condition was performed in triplicate. The cells were kept at 37 °C with 5 % CO_2_ for 4 days to allow spheroid formation. The spheroids were imaged before treatment. On the fifth day, the cells were treated. Four days post-treatment, the cells were imaged again, and the size of the spheroids was measured. High-content screening (HCS) analysis was performed using a Revvity Operetta-CLS system with a 5X NA 0.4 objective. Image acquisition was carried out using Harmony software (Revvity). Image analysis and nucleus counting were performed using Columbus software (Revvity).

### Caspases 3 and 7 Activity

For each condition, cells were seeded in flat-bottom 96-well plates and incubated for 24 hours. The cells were then treated for 15 hours under different conditions: untreated, treated with the rabbit anti-eL21 polyclonal antibody, and treated with a rabbit polyclonal IgG isotype control. Each condition was performed in triplicate. To measure caspase activity, 100 *µ*L of Caspase-Glo 3/7 reagent (Promega, G8090) was added to the cells for 30 minutes at room temperature, protected from light. Fluorescence intensity was measured using the Tecan I-Control system at 570 nm.

### Cell Cycle

For each condition, cells were seeded in flat-bottom 96-well plates and incubated for 24 hours. The cells were then treated for 15 hours under different conditions: untreated, treated with the rabbit anti-eL21 polyclonal antibody, treated with nocodazole (Sigma-Aldrich, M1404), and treated with thymidine (Sigma-Aldrich, T1895-5G). Each condition was performed in triplicate. EdU 1X at 1 mM was added to the appropriate cell culture medium for 60 minutes at 37 °C and 5 % CO_2_. The medium was then removed, and the cells were fixed with PBS 1X containing 4 % PFA for 5 minutes at room temperature. The cells were permeabilized with PBS 1X containing 0.5 % Triton X-100 for 20 minutes. The cells were then incubated in PBS 1X containing 2 mM CuSO_4_.H_2_O, 4 *µ*M Sulfo-Cy^3^-Azide, and 20 mg/mL ascorbic acid for 30 minutes at room temperature, protected from light, and sealed with parafilm to prevent oxidation. Cells were saturated in antibody buffer containing PBS 1X, 1 % BSA, 2.2 % NaCl, and 0.5 % Tween 20 for 2 hours. The cells were incubated for 1h with a rabbit polyclonal anti-phospho histone H3 (Ser10) antibody (Cell Signaling, 9701S), and then detected using the Alexa Fluor-555 goat anti-rabbit secondary antibody (Life Technologies, A21428) at a 1:1,000 dilution. Nuclei were stained with Hoechst at a 1:10,000 dilution in PBS 1X. All washing steps were performed using PBS 1X containing 10 mM glycine. High-content analysis was used to measure each fluorescent signal and compute the number of cells in each phase of the cell cycle, using a Revvity Opera Phenix Plus with a 20X NA 0.4 objective. Image acquisition was carried out using Harmony software (Revvity). Image analysis and nucleus counting were performed using Columbus software (Revvity).

### Statistics

Statistical analyses were performed using GraphPad Prism 10 software. T-test (unpaired, non-parametric was used for statistical significance.

## Results

### Detection of eL21 on the surface of triple-negative breast cancer cells

Based on the externalization of uL10 in tumor cells [28], [29], we hypothesized that other ribosomal proteins might be abnormally localized at the surface of cancer cells. Using subcellular fractionation on TNBC cell lines CAL51 and Hs 578T, we tested for the presence of various ribosomal proteins in the plasma membrane fraction (Fig. 1A). The subcellular fractionation revealed the presence of uL18 (RPL18) and eL21 in the plasma membrane fraction in both cell lines. In contrast, uL10, eL19 (RPL19) and uS5 (RPS2) were not found in the plasma membrane fraction of these cell lines, despite a similar abundance in the cytoplasmic fraction. The membrane-associated form of eL21 is present in both TNBC cell lines CAL51 and Hs 578T, with expression approximately 3 times higher in CAL51. (Fig. 1B). We pursued the study with eL21, by evaluating its presence at the surface of the cells by immunofluorescence. Staining was performed with a polyclonal anti-eL21 antibody, raised against the 110-160 domain, on both permeabilized cells (Fig. 1C upper panel) and non-permeabilized cells (Fig. 1C lower panel) to analyze the presence of eL21 (green) in the intracellular compartment and at the cell surface, respectively. Detection of Na^+^K^+^ATPase (red), which is sensitive to permeabilizing agents, was used as a marker of plasma membrane. In permeabilized cells, eL21 was homogeneously present throughout the cytoplasm, as expected for a ribosomal protein (Fig. 1C). Interestingly, in non-permeabilized cells, fluorescent spots were observed in both cell lines (Fig. 1C). The staining pattern of eL21 was markedly different between permeabilized and non-permeabilized cells, indicating that the signal observed in non-permeabilized cells, was not due to unwanted detection of intracellular eL21. eL21 was detected at the surface of most but not all cells. Furthermore, we noticed that in CAL51 and Hs 578T cells positive for eL21 signal were also positive for Na,K ATPase signal.

**Figure 1.**
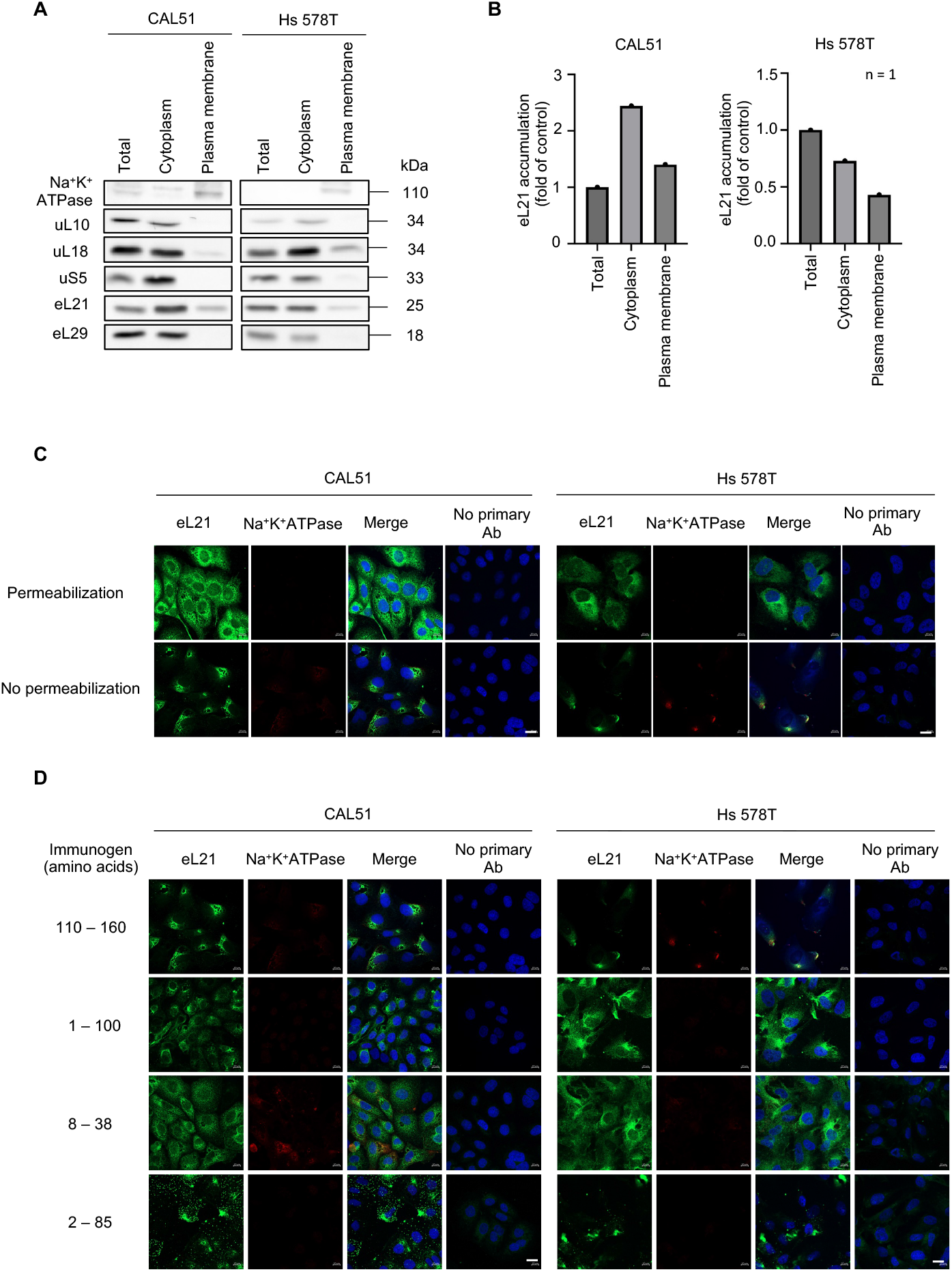
Evidence of eL21 at the surface of TNBC cells. (**A**) Western blot detection of ribosomal proteins upon subcellular fractionation and plasma membrane purification. Na^+^K^+^ATPase was used as plasma membrane marker. (**B**) Quantification of eL21 in each fraction shown in A. Data are normalized to the total lane. (**C**) Immunofluorescence of eL21 on either permeabilized or non-permeabilized cells. eL21 was detected with antibody raised against the 110-160 region (Supp. Fig. 1). Na^+^K^+^ATPase is used as a marker of cell permeabilization. Nuclei were stained with Hoechst. Scale = 20 μm. (**D**) Immunofluorescence performed on non-permeabilized cells. The region recognized by each eL21 antibody is indicated on the left. Na^+^K^+^ATPase is used as a marker of cell permeabilization. Nuclei were stained with Hoechst. Scale = 20 μm. Data are representative of at least two independent biological replicates.

To strengthen this observation, we performed immunofluorescence on TNBC cells with additional anti-eL21 antibodies generated against different parts of eL21, spanning the entire protein (Fig. 1D and Supp Fig. 1A). Fluorescence was observed on non-permeabilized cells with all four antibodies and in both cell lines, thus consolidating the evidence of a cell surface form of eL21. Moreover, since the four antibodies recognize epitopes spread all along eL21 (Supplementary Fig. 1A), the data suggests that the entire eL21 could be present at the surface. Notably, both immunofluorescence and western blot data are consistent with the presence of the full length eL21 at the surface of the cells (Fig.1A and 1D, and Supplementary Fig. 1B).

To consolidate our data, the presence of eL21 at the cell surface was then evaluated using transmission electron microscopy analysis on ultrathin sections of CAL51 and Hs 578T cells (Fig. 2A). Staining was performed using the antibody directed against the C-Terminal part of eL21 (Fig. 2A) and compared to EGFR staining as a positive control of a membrane protein (Fig. 2A). eL21 was found in the cytoplasm as expected for a ribosomal protein. In addition, eL21 was detected at the membrane either alone or in clusters (indicated by red triangles). As expected, EGFR signal was observed only at the plasma membrane. Electron microscopy allows the distinction between extracellular and internal side of the cell, using the distance between the cell membrane and the gold particle. Based on the size of an antibody which define the maximum distance of the protein of interest from the gold particle, we quantified the presence of eL21 in membrane or in the internal side of the plasma membrane versus on the extracellular leaflet of the membrane, in each image (Fig. 2B). Among the membrane associated signal over 95% of the gold particles were in the category “intramembranous or extracellular” which was similar to the signal of EGFR (Fig. 2B). Thus this result further support that eL21 is localized on the extracellular leaflet of the membrane (Fig. 2B).

**Figure 2.**
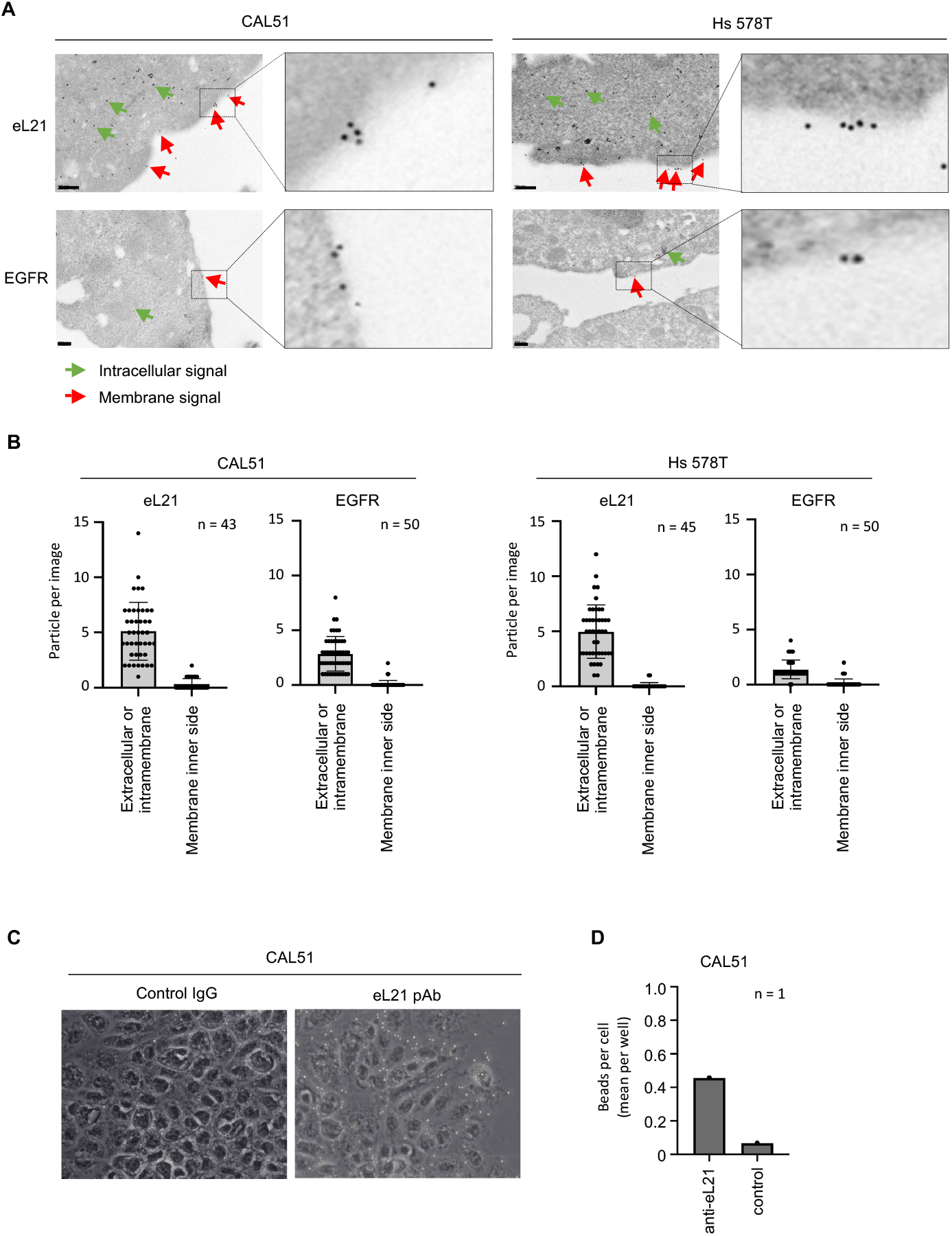
eL21 is present at cell surface of TNBC cell lines. (**A**) Immunogold detection of eL21 and EGFR proteins on ultrathin section of CAL51 and Hs578T cells by transmission electronic microscopy analysis. Intracellular and membrane associated signals are indicated with arrows. Data are representative of at least 40 images. Scale bar is 200 nm. (**B**) Quantification of the position of gold conjugated secondary antibody with regard to the plasma membranes. The distance with membrane was used to discriminate between intramembranous / extracellular signal and intracellular signal. The data are counting from at least 40 different images (**C**) Microbeads binding assay. Phase-contrast images of non-permeabilized cells upon immune-labeling with eL21 antibody and detection with protein-G coupled microbeads. Assay was realized with eL21 antibody binding to the C-terminal region of eL21 (Supplementary Fig. 1A) Controls were performed without primary antibody. (**D**) Quantification of microbeads number from the experiment in (**C**). Image analysis was used to count the beads and nuclei.

The experiments presented above, strongly support the presence of eL21 at the cell surface. However, the resolution of confocal microscopy and immuno-detection by electron microscopy prevents from unquestionably demonstrating that the protein is at the extracellular side of the cell. To firmly demonstrate that the eL21 is present at the surface of the cell, we set-up an approach to detect the attachment of eL21 antibodies on non-permeabilized cells, using protein G coupled microbeads, which cannot cross the cell membrane due to their size (2.8 **µ**m) (Fig. 2C-D). The primary antibody directed against the C-Terminal part of eL21 was incubated with non-permeabilized cells, followed by incubation of the cells with the protein G coupled microbeads. The beads bound to the cells appeared as refringent white dots on the phase contrast images, allowing for their visualization and quantification (Fig. 2C). Clusters of beads were observed on the surface of both cell lines, indicating the presence of an extracellular form of eL21 (Fig. 2C and 2D). As a negative control, the binding of protein G coupled microbeads was monitored without primary antibody incubation (Fig. 2C). The presence of microbeads bound at the plasma membrane was found to be approximately 6.7 times higher in CAL51 compared to the negative control without the primary antibody (Fig. 2D). In conclusion, these results demonstrate the presence of a novel form eL21 on the surface of TNBC cell lines.

### Immunotargeting of the cell surface form eL21 induces a strong cytotoxic effect in triple-negative breast cancer cell lines

Cell surface targets can be used to lead a toxic payload into tumor cells but also to trigger a cytotoxic effect as a direct consequence of antibody binding. We evaluated the impact of targeting the externalized form of eL21 with the polyclonal anti-eL21 antibody raised against the 8-38 region of eL21. We first evaluated cell proliferation by monitoring DNA replication using an EdU incorporation. A strong loss of DNA replicating cells was observed in both cell lines treated 96h with the anti-eL21 antibody at 5 *µ*g/mL, with a 95 % and 92 % reduction in EdU-positive nuclei compared to untreated cells for CAL51 and Hs 578T, respectively (Fig 3A). Additionally, we monitored cell proliferation in real time (Fig. 3B). Treating cells with the anti-eL21 antibody induced a complete block in cell proliferation of both cell lines, compared to cells treated with a control IgG (Fig 3B). To exclude any off-target effect due to non-specific binding of the primary antibody, the anti-eL21 antibody was competed with full-length recombinant eL21 protein, using increasing eL21:anti-eL21 molarity ratios. While low ratios of 0.5:1 and 1:1 had no impact, 5:1 and 10:1 ratio, recombinant eL21 clearly alleviated the growth inhibition effect of the anti-eL21 antibody (Fig. 3C). At 112 hours post-treatment, at the 10:1 ratio, cell growth was restored at values close to that of untreated cells, with XX % and XX % for CAL51 and Hs 578T respectively, compared 73 % and 99 %, respectively for non-treated cells. This indicated that the effect of the anti-eL21 antibody on cell proliferation is due to the specific binding the antibody to cell surface eL21. Next, we evaluated the dose-dependent effect of the anti-eL21 antibody. To this end, cell confluence was monitored with untreated cells compared to those treated with anti-eL21 antibody, ranging from 0.5 to 5 *µ*g/mL. For both CAL51 and Hs 578T, the anti-eL21 antibody induced a dose-dependent inhibition of cell proliferation, ranging from 6 % for 5 *µ*g/mL to 69 % for 0.5 *µ*g/mL for CAL51, and from 9 % for 5 *µ*g/mL to 64 % for 0.5 *µ*g/mL for Hs 578T (Fig. 3D). Based on these results, the anti-eL21 antibody was used at 5 *µ*g/mL in the following experiments.

**Figure 3.**
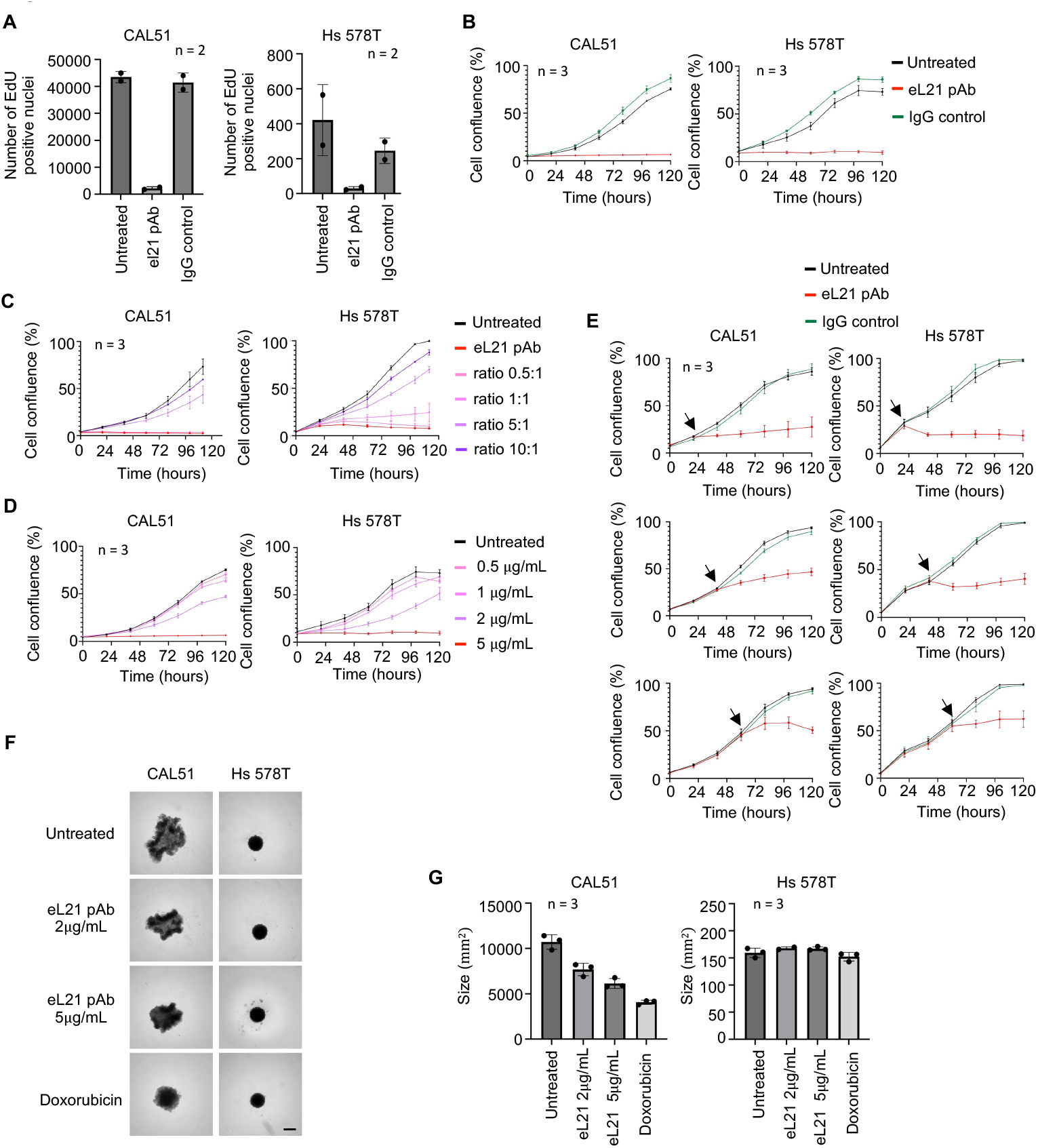
Targeting externalized eL21 reduces cell proliferation of TNBC cells. (**A**) Cellular proliferation assay using EdU incorporation. Cells were either untreated or treated with eL21 pAb (HPA047252) at 5 μg/mL or IgG isotype control at 5 μg/mL for 96 hrs. Data are mean values +/-sd of two independent replicates (**B**) Real-time imaging cell proliferation assay. Cells were either untreated or treated with eL21 pAb (HPA047252) at 5 μg/mL or IgG isotype control at 5 μg/mL. Data are mean values +/-sd of 3 technical replicates. Data is representative of three independent replicates (**C**) Competition assay of eL21 antibody with eL21 purified protein. Cell growth was monitored using real-time imaging cell proliferation assay. Cells were either untreated or treated with eL21 pAb (HPA047252) at 5 μg/mL alone or in combination with human recombinant eL21 protein at the indicated molar ratio. Data are mean values +/-sd of 3 technical replicates. Data is representative of three independent replicates. (**D**) Dose response assay of eL21 antibody. Cell growth was monitored using real-time imaging cell proliferation assay. Cells were either untreated or treated with eL21 pAb (HPA047252) at at the indicated concentration. Data are mean values +/-sd of 3 technical replicates. Data is representative of three independent replicates. (**E**) Cell growth was monitored using real-time imaging cell proliferation assay. Cells were either untreated or treated with eL21 pAb (HPA047252) at 5 μg/mL, 24, 48 or 72 hours after seeding. Data are mean values +/-sd of 3 technical replicates. Data is representative of three independent replicates. (**F**) Spheroids growth assay. Spheroids were allowed to grow for 4 days, and then left untreated or treated with eL21 pAb (HPA047252) at 2 μg/mL or IgG isotype control at 2 μg/mL or Doxorubicin at 2 μM. Each condition was performed in triplicate. (**G**) Spheroids size measurement of the experiment in (**F**) using image analysis. Data are mean values +/-sd of 3 technical replicates. Data is representative of three independent replicates.

In the above experiments, we noted that cell proliferation was inhibited from the early time points, raising the possibility that the L21 antibody might interfere with cell attachment to the well surface. To exclude any interference with cell adhesion, cells were treated after cell adhesion and spreading, at different time points (Fig. 3E). At 24, 48, and 72 hours after seeding, while cells had started proliferating, the eL21 antibody induced a proliferation arrest (Fig. 3E), similar to that observed when treatment was applied at seeding. Notably, the proliferation arrest was observed less than 2 hours after treatment.

Finally, the impact of eL21 antibody was studied on both CAL51 and Hs 578T cultured in 3D as spheroids (Fig. 3F), and compared to doxorubicin, one of the first line chemotherapeutic drugs in TNBC (Fig. 3F). The anti-eL21 antibody had a clear growth inhibitory effect on CAL51 spheroids, with a 29 % reduction in spheroid size after 4 days of treatment at 2 *µ*g/mL and a 43 % reduction at 5 *µ*g/mL compared to untreated spheroids (Fig. 3F-G). However, no significant effect was observed on Hs 578T spheroids both with anti-eL21 antibodies and doxorubicin (Fig. 3F-G). Together, these results showed that targeting cell surface eL21 with an anti-eL21 antibody strongly and rapidly inhibits cell proliferation.

### Targeting of externalized eL21 induces apoptosis and disrupts the cell cycle of TNBC cells

To determine the underlying mechanism of cytotoxic effect of the anti-eL21 binding to cell surface eL21 we searched for cell cycle alteration and cell death by apoptosis. The loss of DNA replicating cells upon 96h of treatment (Fig. 3A) indicated that cells did not progress through the cell cycle. The impact of targeting eL21 on the cell cycle was studied after 15 hours of treatment with the anti-eL21 antibody on both cell lines (Fig. 4A). CAL51 cells treated with the anti-eL21 antibody showed a decrease in the S phase and an increase in the G1 and G2/M phases, suggesting a blockade both at the G1/S and G2/M transition. For Hs 578T cells, treatment with the anti-eL21 antibody resulted in a decrease in the S phase with an increase in the G1 phase, suggesting a blockade in the G1/S transition.

**Figure 4.**
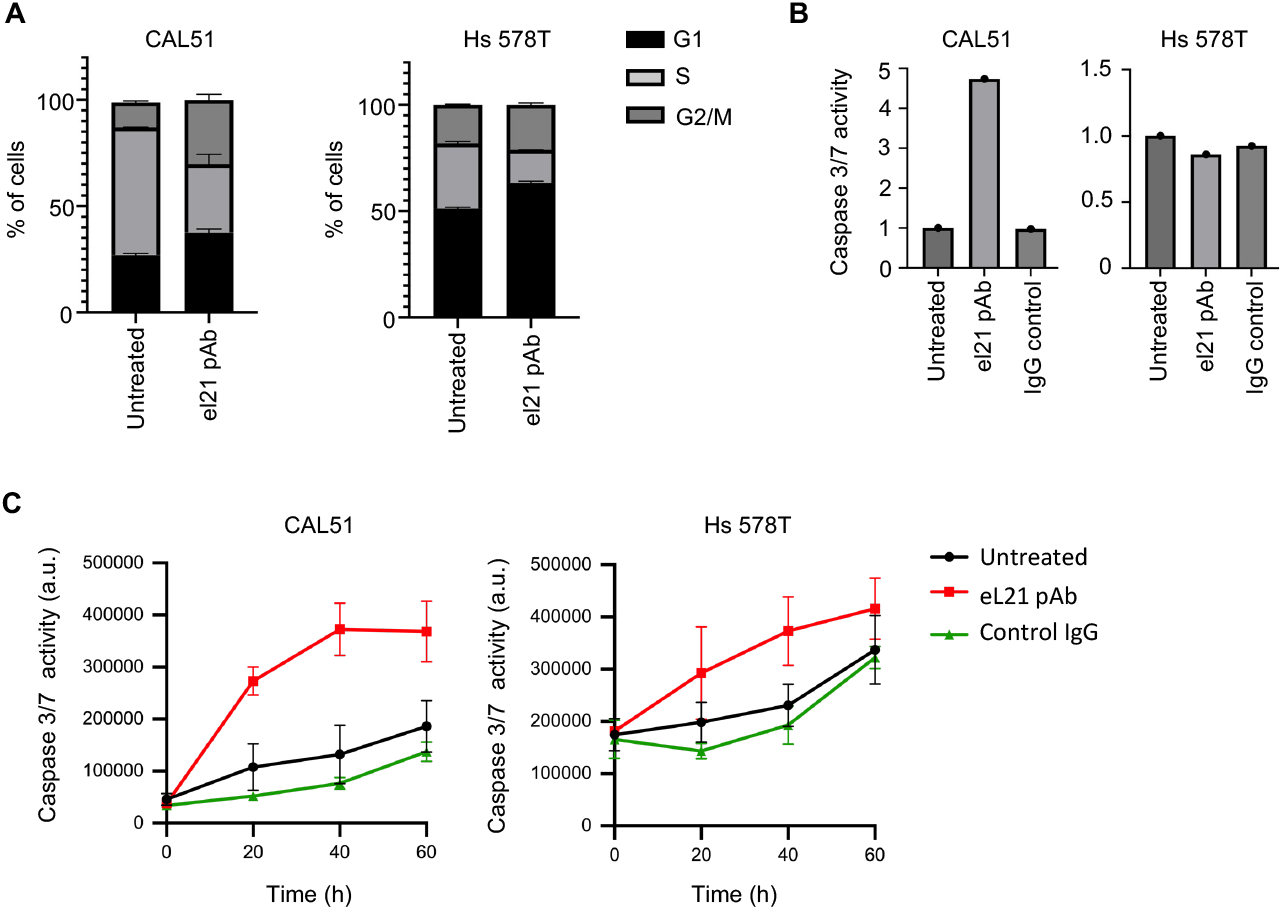
Targeting of externalized eL21 induces apoptosis and disrupts the cell cycle of TNBC cells. (**A**) Cell cycle analysis. Cells were either untreated or treated with eL21 pAb (HPA047252) at 5 μg/mL for 15 hours. High-content analysis was used to measure EdU incorporation and Hoechst nuclear staining. Data are mean values +/-sd of 3 technical replicates. (**B**) Caspase 3/7 activity apoptosis assay. Cells were either untreated or treated with eL21 pAb (HPA047252) at 5 μg/mL or IgG isotype control at 5 μg/mL for 15 hrs. Data were normalized to the untreated condition. (**C**) Kinetic Caspase 3/7 apoptosis assay. Cells were either untreated or treated with eL21 pAb (HPA047252) at 5 μg/mL or IgG isotype control at 5 μg/mL for 60h. Data are mean values +/-sd of 3 technical replicates.

Next, we analyzed apoptosis by monitoring the activation of effector caspases 3 and 7. Initially, the activation of caspases 3 and 7 was monitored after 15 hours of treatment with the anti-eL21 antibody (Fig. 4B). A strong activation of caspases was observed in CAL51 cells treated with eL21 antibody compared to the IgG control. In contrast, no significant caspase activation was observed in Hs 578T cells. To evaluate whether a longer treatment than 15 hours was necessary to induce apoptosis, we monitored Caspase 3/7 activation by kinetic analysis over 72 hours of treatment (Fig. 4C). The activation of caspases 3/7 in CAL51 cells treated with the anti-eL21 antibody was observed less than 2 hours post-treatment and reached 4.7-fold the signal of untreated cells. For Hs 578T cells, a non-significant increase in caspase activation was observed, with a signal 0.8 times higher than in untreated cells. Taken together, these results indicate that targeting eL21 on the surface of TNBC cells disrupts the cell cycle and activates an apoptosis response in a cell type dependent manner.

## Discussion

In this study, we uncovered the presence of the ribosomal protein eL21 on the surface of TNBC cells, and we make the proof of concept that using an anti-eL21 antibody, it is possible to target the cell surface eL21 and induce a strong anti-proliferative effect and activate apoptosis.

Our study expands the list of proteins that are externalized in cancer cells, by adding a second RP. Externalized proteins, have significant therapeutic interest, since their presence on the surface of cancer cells provides an opportunity to exploit antibody-based or ligand-based strategies. For instance, the proof of concept that externalized protein are actionable was made with the CK8 protein, a cytoskeleton protein externalized on the surface of colorectal cancer cells and of poorly differentiated head and neck squamous cell carcinoma cells [13], [22]. In addition, a role in invasion has been described for externalized CK8, making it an interesting therapeutic target for metastatic cancers [23], [22].

Extracellular proteins are characterized by the lack of conventional membrane localization signal that may explain their localization. Consequently, it is necessary to carefully demonstrate their unexpected localization at the surface of the cell. Here, we demonstrate the existence of an extracellular form of eL21 in TNBC cells using multiple complementary cell biology and molecular approaches. Immunofluorescence analysis revealed the extracellular form of eL21 as discrete spots, while its intracellular form was distributed homogeneously throughout the cytoplasm. Subcellular fractionation confirmed the association of eL21 with the plasma membrane, eL21 being detected in membrane fraction. Electron microscopy further validated this finding, showing eL21 on the extracellular side of the membrane, either as individual spots or in clusters. Finally, an assay based on the binding of protein G microbeads on the surface of cells confirmed the extracellular accessibility of eL21. This observation has two key implications: first, unlike antibodies, which may pass through damaged plasma membrane, microbeads cannot penetrate cells, firmly demonstrating the presence of eL21 on the surface of TNBC cells. Second, it demonstrates that eL21 is accessible to antibody present in the extracellular environment, highlighting its potential in antibody-based therapies. Importantly, multiple antibodies were used to consolidate the presence of eL21 on the cell surface, including three polyclonal and one monoclonal antibody targeting eL21 peptides spanning the full 160 amino acids of the protein. Western blot analysis of the plasma membrane fraction detected eL21 at its full length. Together, these findings strongly suggest that the complete eL21 protein is externalized in TNBC cells.

The presence of a ribosomal protein, uL10, on the surface of breast cancer, head and neck cancer and colorectal cancer cells has recently been described [28], [29]. uL10 appears to be exposed on the cell surface during specific cellular processes, involving inflammation and cellular stress [28], [29]. In contrast, eL21 was found externalized on normally growing TNBC cells. While this indicated that eL21 externalization does not rely on cell stress, it is possible that cellular stress modulates eL21 externalization. Exposing TNBC cells to eL21 antibodies induced cell cycle arrest and triggered the apoptotic pathway, a phenomenon also observed with anti-uL10 antibodies [28]. These observations highlight yet unknown extra-ribosomal functions of eL21 and uL10. Cell surface functions of these RP would add to the list of already known extra-ribosomal functions of ribosomal proteins, including, free uL1 (RPL10A) involved in the transcriptional regulation of certain genes in plants [30], or eS6 (RPS6), which plays a role in ribosomal DNA transcription [31], uS3 (RPS3) which is involved in DNA repair, and uS11 (RPS14), which participates in the autogenous regulation of replication. In this context, externalized eL21 may modulate cell behavior by communicating to the intracellular compartment, information regarding the extracellular environment. Notably, anti-uL10 antibodies have been found in the sera of breast cancer and colorectal cancer patient [29], which underscores the status of externalized RP as tumor-specific antigens, and raised the question of an immune-response against externalized eL21 and breast cancer patients.

We find that exposing TNBC cells to anti-eL21 antibodies strongly impacted cell proliferation. The anti-eL21 antibody, generated against the N-terminal part of the eL21 protein, showed a dose-dependent effect on cell proliferation, inducing a complete halt in cell proliferation, regardless of the timing of treatment. We provide two possible mechanisms to explain this cessation of proliferation: first a cell cycle arrest, and second, activation of apoptosis. While cell cycle was significantly impacted in both TNBC cell lines used in this study, significant activation of effector caspases 3 and 7 was observed only in CAL51 cells and not in HS578T cells, the latter being described as more resistant to apoptosis. Indeed, several breast cancer cell lines with p53 mutations have been described as more resistant to apoptosis, as is the case for HS578T, which has a p53 missense mutation VI57F resulting in loss of p53 function [34], [35], [36]. Furthermore, Hs 578T cells overexpress the genes Blc-2 and Bcl-xL, which represses intrinsic apoptosis pathways [37], [38], [39]. Other potential mechanism of action of anti-eL21 antibody may include the internalization of the eL21 antibody and perturbation of intracellular eL21 activity in ribosome assembly and activity in translation. Indeed, eL21 down-regulation by siRNA in HeLa cells inhibits ribosome maturation, characterized by an accumulation of 12S and 32S pre-ribosomal RNAs and activation of the p53, and induction of a strong ribosomal stress response [42]. In addition, preliminary studies have investigated the role of eL21 in pancreatic cancer. siRNA-based targeting of eL21 in pancreatic cancer cells appears to inhibit cell proliferation and DNA replication, inducing cell cycle arrest and apoptosis [33]. While this observation may result from depletion in intracellular eL21, the toxic effect of eL21 knock-down may also be explained by the loss of extracellular-function of eL21.

Taken together, these findings identify eL21 as a novel target on the surface of TNBC cells, eL21, where its activation by an antibody elicits a strong antitumoral effect. Additionally, they support the existence of a previously unrecognized extra-ribosomal function of ribosomal proteins at the cell surface. Given its localization and the cytotoxic response upon immuno-targeting, eL21 emerges as a promising therapeutic target for TNBC.

## Declarations

## Ethics approval and consent to participate

Not applicable

## Consent for publication

Not applicable

## Availability of data and material

All raw data generated during the study are available from the corresponding authors upon request. Biological material is available upon reasonable request.

## Authors contributions

LA performed experiments and analysed data. NJ performed experiments. NRB, FPL, MAA and FC conceived the study. FC analysed data and supervised the study. JJD and FC secured funding. LA and FC wrote the manuscript. JJD, MAA and FC reviewed the manuscript.

## Funding

The study was supported by grants from the Canceropole Lyon Auvergne Rhone Alpes to FC (CLARA, Oncostarter, RibExt Program), Agence Nationale de la Recherche (LabEx DEVweCAN, ANR-10-LABX-0061; Institut Convergence ANR-17-CONV-0002), European network COST Translacore (COST Action CA21154).

F.C. is a CNRS research fellow. J.-J.D. is an Inserm research fellow.

## Aknowledgement

We thank Christophe Vanbelle and Mélina GAUTIER (Cellular Imaging Platform of the Cancer Research Center) for technical help. We thank Elisabeth Errazuriz-Cerda (CIQLE Platform) for help with electron microscopy analysis. We thank Brigitte Manship for editing the manuscript.

