## Supplementary Figures for "Identification of ribosomal protein eL21 as a novel externalized protein and a potential target in triple negative breast cancer"

**A**

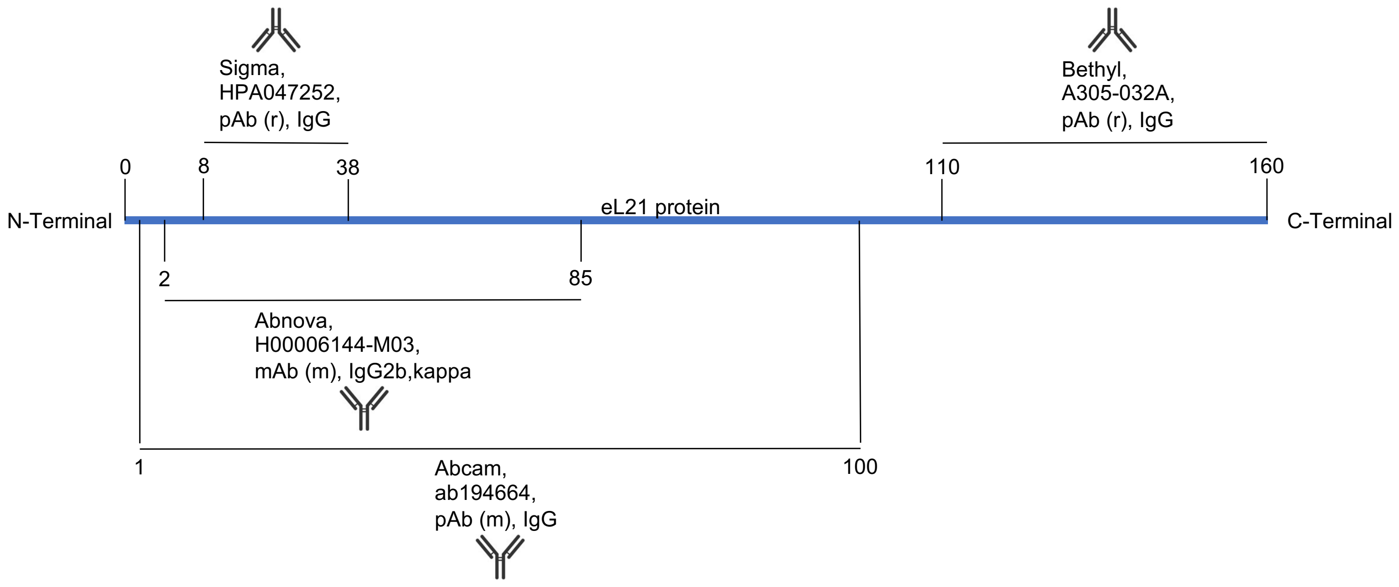


**B**

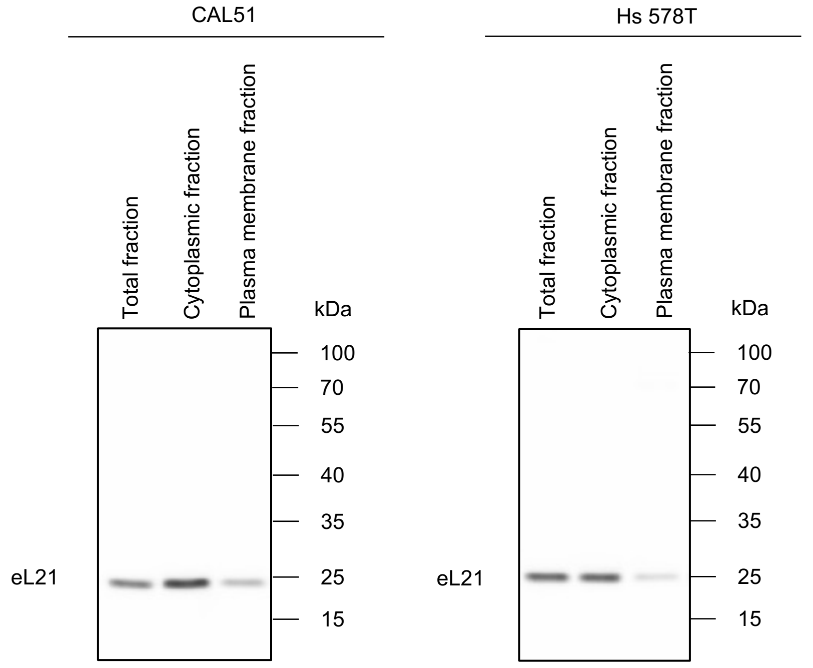


**Supplementary Figure 1.**
**A**. Schematic representation of eL21 protein (blue line, 160 AA) with the immunogen used to raise the anti-eL21 antibodies used in the study.
**B**. Uncropped western blot images of eL21 detection from Figure 1A. The image shows that eL21 is detected only as a full size 25 kDa form.
